# The CFDE Workbench: Integrating Metadata and Processed Data from Common Fund Programs

**DOI:** 10.1101/2025.02.04.636535

**Authors:** John Erol Evangelista, Daniel J.B. Clarke, Zhuorui Xie, Stephanie Olaiya, Heesu Kim, Giacomo B. Marino, Anna Byrd, Shivaramakrishna Srinivasan, Sumana Srinivasan, Mano R. Maurya, Sherry L. Jenkins, Andrew D. Lutsky, Lucas Sasaya, Alexander Lachmann, Nasheath Ahmed, Ido Diamant, Ethan Sanchez, Srinivasan Ramachandran, Shankar Subramaniam, Avi Ma’ayan

## Abstract

The NIH Common Fund Data Ecosystem (CFDE) program was established to facilitate data accessibility and interoperability across multiple Common Fund (CF) programs and promote collaborations and new discoveries by combining data from different CF programs. The CFDE Data Resource Center (DRC) was tasked with developing two web-based portals: an Information Portal to serve information about the CFDE, and a Data Portal to host harmonized metadata and processed data contributed by participating CF Data Coordination Centers (DCCs) and other sources. To achieve these goals, the CFDE DRC developed the CFDE Workbench, a web-based platform that hosts highly processed data, metadata, tools, use cases, and analyses developed by the CFDE. The Crosscut Metadata Model (C2M2) and several other processed data formats are hosted by the CFDE Workbench, including set libraries (XMTs), Knowledge Graph (KG) assertions, and attribute tables. These processed data formats make information derived from CF programs more findable, accessible, interoperable, and reusable (FAIR) and artificial intelligence (AI)-ready for cross-DCC knowledge discovery. In addition to these processed data formats, several other assets are served by the Workbench, including Extract Transform Load (ETL) pipelines, OpenAPI-documented and SmartAPI-registered Representational State Transfer (REST) Application Programming Interfaces (APIs), entity pages such as gene, drug, disease, or tissue landing pages, Playbook Workflow Builder (PWB) metanodes which are codified workflow steps involving DCC APIs, and chatbot specifications which are prompts helping chatbots to use DCC APIs to answer a wide range of user queries. Besides serving data, metadata, and code assets, the CFDE Workbench also has several tools that use these data, metadata, and code assets to enable cross-CF-program knowledge discovery use cases. Overall, the CFDE Workbench is a platform that consolidates efforts toward making CF programs funded resources harmonized, FAIR, and AI-ready. The CFDE Workbench website is available from: https://cfde.info/.

## INTRODUCTION

### The Common Fund Data Ecosystem (CFDE) and the Data Resource Center (DRC)

Since its establishment in 2006, the National Institutes of Health (NIH) Common Fund (CF) has supported significant advancements in biomedical research by funding ambitious, cross-institutional initiatives. Examples of CF programs that are making a big impact on the biomedical research community include the Genotype-Tissue Expression (GTEx) [1], the Human Microbiome Project (HMP) [2], Undiagnosed Diseases Network (UDN) [3], Knockout Mouse Phenotyping Program (KOMP2) [4], Metabolomics [5], Glycoscience [6], Kids First [7], Human Biomolecular Atlas Program (HuBMAP) [8], Stimulating Peripheral Activity to Relieve Conditions (SPARC) [9], Illuminating the Druggable Genome (IDG) [10], the Library of Integrated Network-Based Cellular Signatures (LINCS) [11], Bridge to Artificial Intelligence (Bridge2AI) [12–15], Cellular Senescence Network (SenNet) [16], Extracellular RNA Communication (ExRNA) [17], and the Molecular Transducers of Physical Activity Consortium (MoTrPAC) [18]. CF programs have produced diverse high-dimensional massive datasets ranging from omics profiling of human and mouse cells and tissues as well as other deep phenotyping data, including imaging and clinical parameters. Many of these datasets are served via online databases and tools that have been widely used by the biomedical research community. Recently, the CF established a new program called the Common Fund Data Ecosystem (CFDE) with the aim at furthering the CF datasets’ findability, accessibility, interoperability, and reusability (FAIRness) [19] and artificial intelligence (AI)-readiness. The goal of the CFDE program is to assist biomedical researchers in forming complex hypotheses that utilize and integrate data generated from several CF programs. The CFDE consortium is made of five centers: the Data Resource Center (DRC), the Knowledge Center (KC), the Integration and Coordination Center (ICC), the Training Center (TC), and the Cloud Working Implementation Center (CWIC). In addition, data coordination centers (DCCs) from participating CF programs are members of the CFDE consortium. Here we describe the primary product of the CFDE Data Resource Center (DRC) which is called the CFDE Workbench.

## METHODS AND RESULTS

### The CFDE Workbench Workflow

The CFDE Workbench is organized into two main components, the Data Portal (https://data.cfde.cloud/) and the Information Portal (https://info.cfde.cloud/). The Data Portal of the CFDE Workbench catalogs several types of uniformly processed data and metadata files and other digital objects from each participating DCC. These files and other assets are submitted to the portal by the participating DCCs and can then be downloaded from the CFDE Workbench Data Matrix or searched for via the CFDE Workbench Data Portal search engine. The Information Portal provides relevant information about each DCC and on overarching consortium activities that include training and outreach events, brief descriptions of CFDE partnership projects, and detailed community-established protocols. Here we describe the various components of the CFDE Workbench and how they all come together to produce a valuable and comprehensive resource for the CFDE consortium and biomedical research community at large (Fig. 1).

**Fig. 1.**
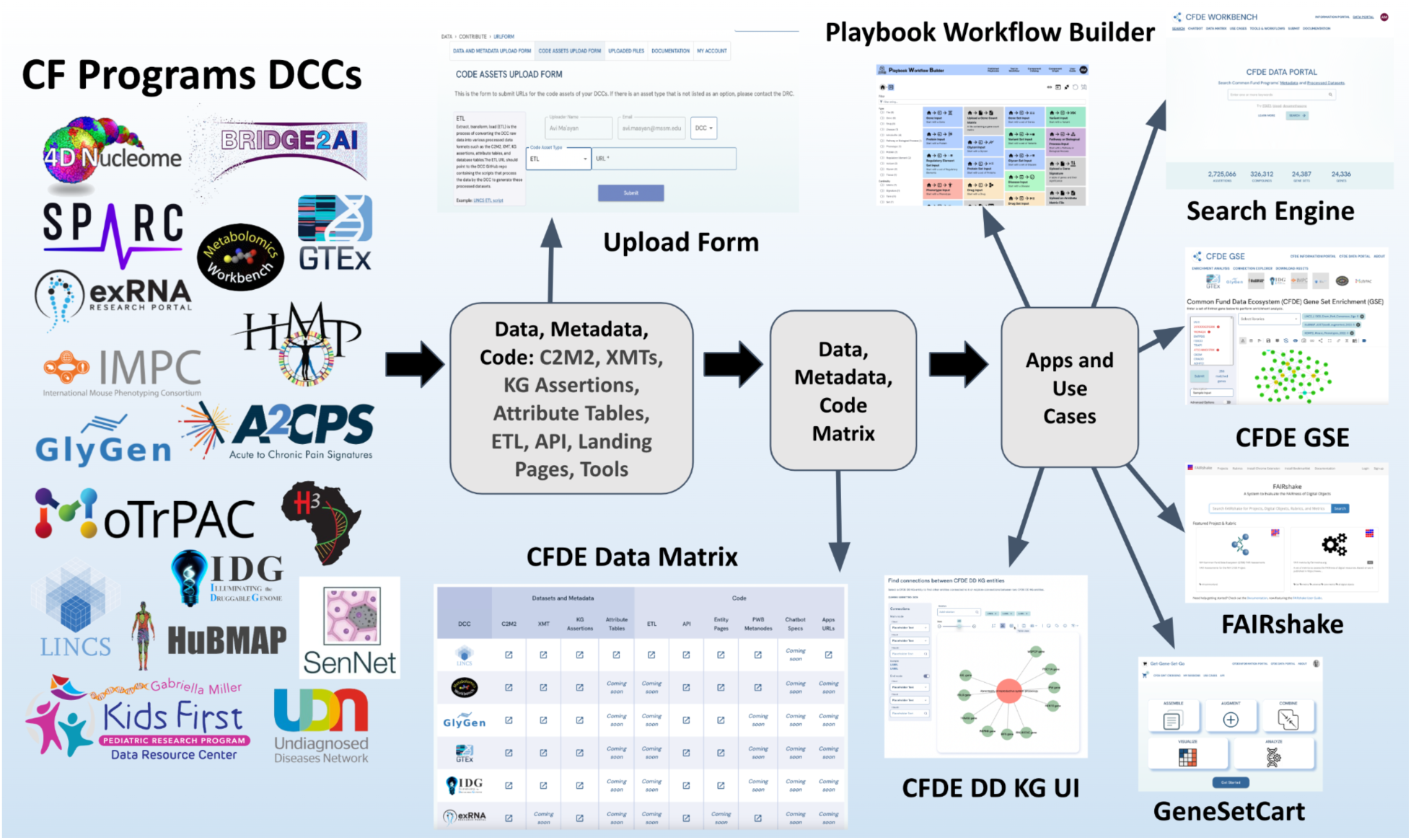
The CFDE Workbench received metadata, processed data, and code assets from the CF program participating DCCs. These assets are uploaded via a data submission system and assessed for FAIRness with FAIRshake. Approved uploads are displayed in a Data Matrix and served via a search engine. The metadata, processed data, and code assets enable the development of a layer of tools that include the PWB, GeneSetCart, DD-KG-UI, and CFDE-GSE.

### The CFDE Workbench Submission System

The CFDE Workbench functions as a hub for finding CF-supported datasets via a dedicated search engine. Digital objects on the CFDE Workbench are broadly grouped into two categories: File assets and code assets. File assets include Cross-Cut Metadata Model (C2M2) [20] packages, knowledge graph (KG) assertions, set libraries (XMT), and attribute tables. Code assets include API documentation links, entity page templates, URLs to apps, extract transpose and load (ETL) scripts, chatbot specifications, and URLs to tools (Table 1). To streamline the collaborative efforts of the CFDE, and to ingest these assets into the CFDE Workbench, the DRC has implemented a data and metadata submission system. This system enables the DCCs to submit their assets to the CFDE Workbench. The data and metadata submission system has two user-friendly input forms: 1) the data and metadata upload form designed for uploading data and metadata files; and 2) the code assets upload form for submitting URLs and descriptions of the various code assets. By using the data and metadata submission system, DCCs can upload and approve their assets for public display on the CFDE Workbench. Upon submission of a new asset, DCC submitters and approvers can use the system to archive their previously published assets and mark their most up-to-date assets as current. In addition, once assets are uploaded, these assets are evaluated for FAIRness with FAIRshake [21] (Fig. 2). Different rubrics have been established to evaluate each asset type. The results from the automated FAIR assessment with FAIRshake are displayed as an insignia near each uploaded asset entry. These FAIRshake insignias are only visible to the data submitters and approvers from each DCC. The FAIR assessment is aimed at giving the DCC submitters and approvers feedback on how they may improve the FAIRness of their uploaded assets. The data and metadata submission system assigns users to one of four roles, each having different access permissions to various pages within the system: a general user, a DCC submitter, a DCC approver, or a DRC reviewer. Comprehensive documentation is provided to assist uploaders and approvers through the process of preparing the assets, submitting them to the workbench, and approving them. The documentation contains details about the asset types, the submission and approval process, as well as descriptions about how to prepare the asset types for publication and search. Overall, by contributing these assets to the workbench, the DCCs can make contributions to the CFDE and make the assets produced by their CF program more findable and accessible.

**Fig. 2.**
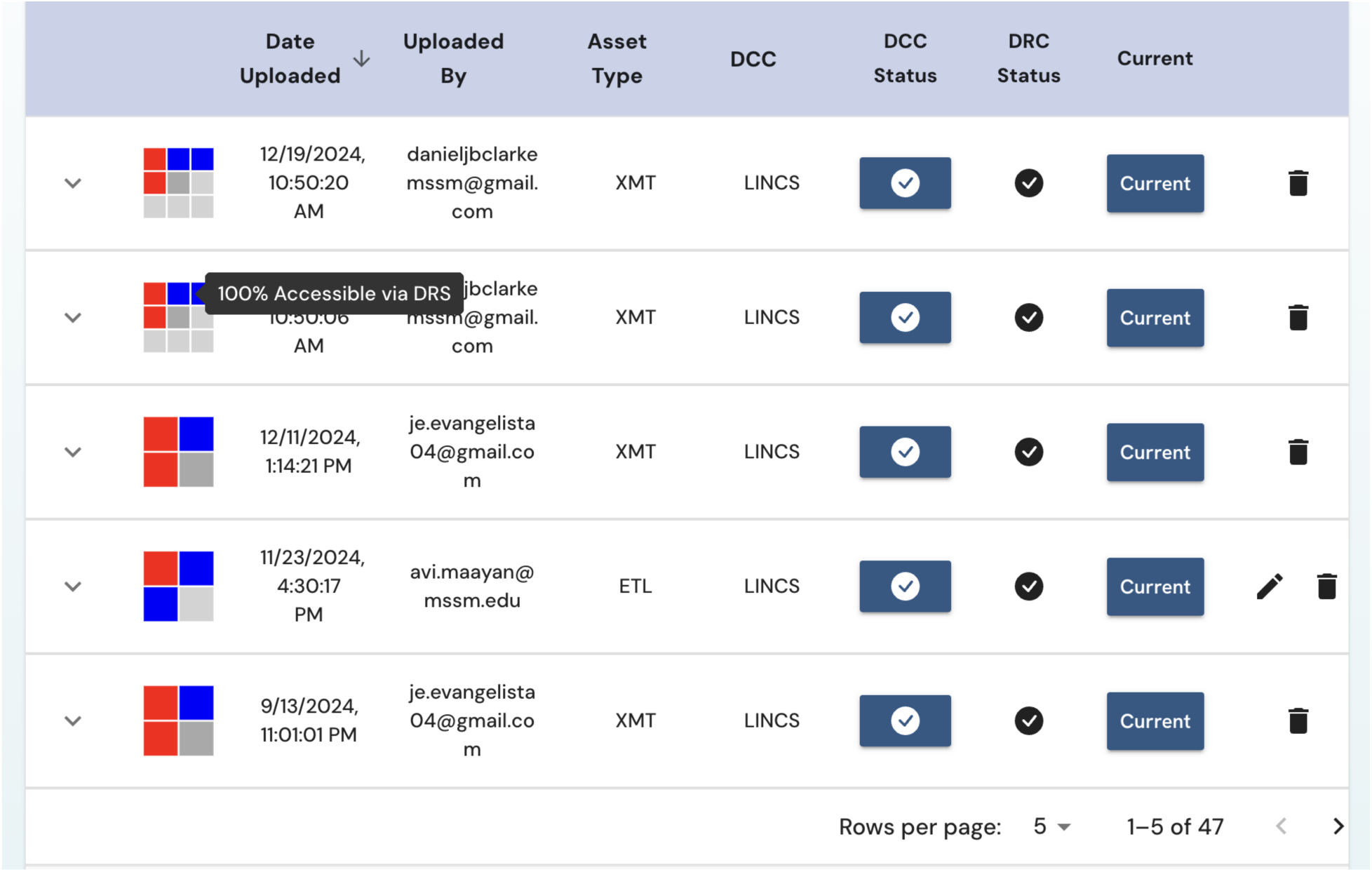
Screenshot from a page within the data submission system that displays a list of assets uploaded by a DCC. Near each uploaded listed asset, the FAIRshake insignia displays the results of the FAIR evaluation of the asset. The table also has interactive switches to approve uploaded assets by the DCCs and the DRC.

**Table 1.** Summary of metadata, processed data, and code assets that are served by the CFDE Workbench and displayed within the Data Matrix.

| Asset type | Data Type | Description | Format |
| --- | --- | --- | --- |
| Datasets and Metadata Assets | C2M2 | A metadata standard for describing data files from a variety of biomedical and biological sources. | zip (tsv) |
|  | XMT | Text based files that contain a collection of sets of an entity type. | gmt or dmt |
|  | KG Assertions | Nodes and edges that describe relationships between different biomedical concepts, objects, and entities. | Zip (csv) |
|  | Attribute Tables | Files containing tables that describe the relationship between two entities with one entity type on the rows (for example genes) and another on the columns (for example tissue types). The table is filled with values that quantify the association between the entities. | h5 or hdf5 |
| Code Assets | ETL | Extract, transform, load (ETL) scripts that process raw data into various processed data formats such as the C2M2, XMT, KG assertions, attribute tables, and database tables. URL redirects to the DCC GitHub repository containing the scripts that process the data by the DCC to generate these processed datasets. | URL, py |
|  | API | REST API to access DCC data and tools. URL redirects to the API documentation. The API recommended standard is OpenAPI, and SmartAPI is the recommended repository for depositing the API docs. | URL |
|  | Entity Pages Template | A template used to create the landing page displaying the datasheet about a gene, a metabolite, and protein, a cell type, or other biological and biomedical entity. | URL |
|  | PWB Metanodes | The components of the PWB that can receive inputs and produce outputs including visualization and data processing operations. URL redirects to a GitHub link that contains the script describing a PWB metanode. | URL (tsx, json) |
|  | Chatbot Specs | The Chatbot Specs enable LLMs to function as specialized chatbots that have access to the exposed API endpoints described in the manifest files and can call these APIs based on user input. URL redirects to a manifest file containing metadata and OpenAPI specifications which can be used to develop a chat plugin for LLMs. | URL |

### Metadata Packages for Capturing Common Metadata Elements

The Cross-Cut Metadata Model (C2M2) packages [20] consist of metadata items organized in approximately 50 related tables. For example, these tables contain records about DCCs, projects, subjects, biosamples, anatomy, disease, collections, and other metadata elements. The use of ontologies and controlled vocabularies is a key feature to ensure the integrity of the C2M2. Some of these ontologies and controlled vocabularies include Uberon [22] for anatomy, and Disease Ontology (DO) [23] for disease. In the C2M2 ingestion and digestion pipeline, as the first step, the metadata from the tab-separated value (tsv) files are ingested into respective database tables stored in a PostgreSQL database. To facilitate meaningful and fast full-text query across all these tables, the CFDE Workbench is utilizing the foreign-key constraints among these tables. These keys are joined to produce a fully flattened (FFL) table that is indexed with respect to multiple columns in the table, such as anatomy, species, disease, assay type, and data type. The FFL table is centered around biosamples since a biosample is the most fine-grained entity in the collection of metadata items across the CF DCCs. Currently, as of January 2025, there is information about ∼2M biosamples stored in the CFDE Workbench. Each biosample comes with additional fields such as anatomy. To incorporate key metadata from files in the search and filtering functions, from the file table which has ∼11M rows, distinct project, assay type and data type fields are saved in another table. This additional table is incorporated into the FFL table. Further, a searchable column, a PostgreSQL tsvector data type, that combines information from all the key columns is added to the FFL table to facilitate the query supporting the search. Similarly, processed data packages such as XMTs and KGs are ingested into their respective tables in the PostgreSQL database, indexed, and further processed to facilitate queries.

### The CFDE Workbench Data Matrix

The CFDE Workbench Data Matrix (https://data.cfde.cloud/matrix) is a key component of the CFDE Workbench Data Portal (Fig. 3). The Data Matrix catalogs available assets from across CF programs. The following 18 CF participating programs contributed, or plan to contribute, to the Data Matrix: 4D Nucleome (4DN) [24,25], Acute to Chronic Pain Signatures (A2CPS), Bridge to Artificial Intelligence (Bridge2AI), Extracellular RNA Communication (ExRNA) [17], Gabriella Miller Kids First Pediatric Research Program (Kids First) [1,25], Genotype-Tissue Expression (GTEx) program [1,26], Glycoscience [6], Human BioMolecular Atlas Program (HuBMAP) [26,27], Human Heredity & Health in Africa (H3Africa), Human Microbiome Project (HMP) [27,28], Illuminating the Druggable Genome (IDG) [10], Knockout Mouse Phenotyping Program (KOMP2) [4], Library of Integrated Network-based Cellular Signatures (LINCS) [11], Metabolomics [5], Molecular Transducers of Physical Activity Consortium (MoTrPAC) [18], Stimulating Peripheral Activity to Relieve Conditions (SPARC) [28], Cellular Senescence Network (SenNet) [29], and Undiagnosed Diseases Network (UDN) [29]. Although individual CF DCCs remain responsible for hosting the original data produced by their specific programs, the CFDE Workbench Data Matrix enables users to access standardized and abstracted representations of knowledge processed from the raw data of different CF programs in one location. The Data Matrix also tracks provenance, performs FAIR assessments on the submitted files, and facilitates DCCs to control asset visibility. The Data Matrix additionally supports integration with external community standards such as the OpenAPI [30] and SmartAPI [31] initiatives, and with internal CFDE efforts to develop original standards such as Playbook Workflow Builder (PWB) metanodes, chatbot specs, and continually extending the C2M2 to support other metadata types. By making this knowledge available for download in community established standardized formats, the CFDE Workbench Data Matrix should be able to catalyze usage and increase the relevance of CF data and metadata assets within the CFDE and beyond.

**Fig. 3.**
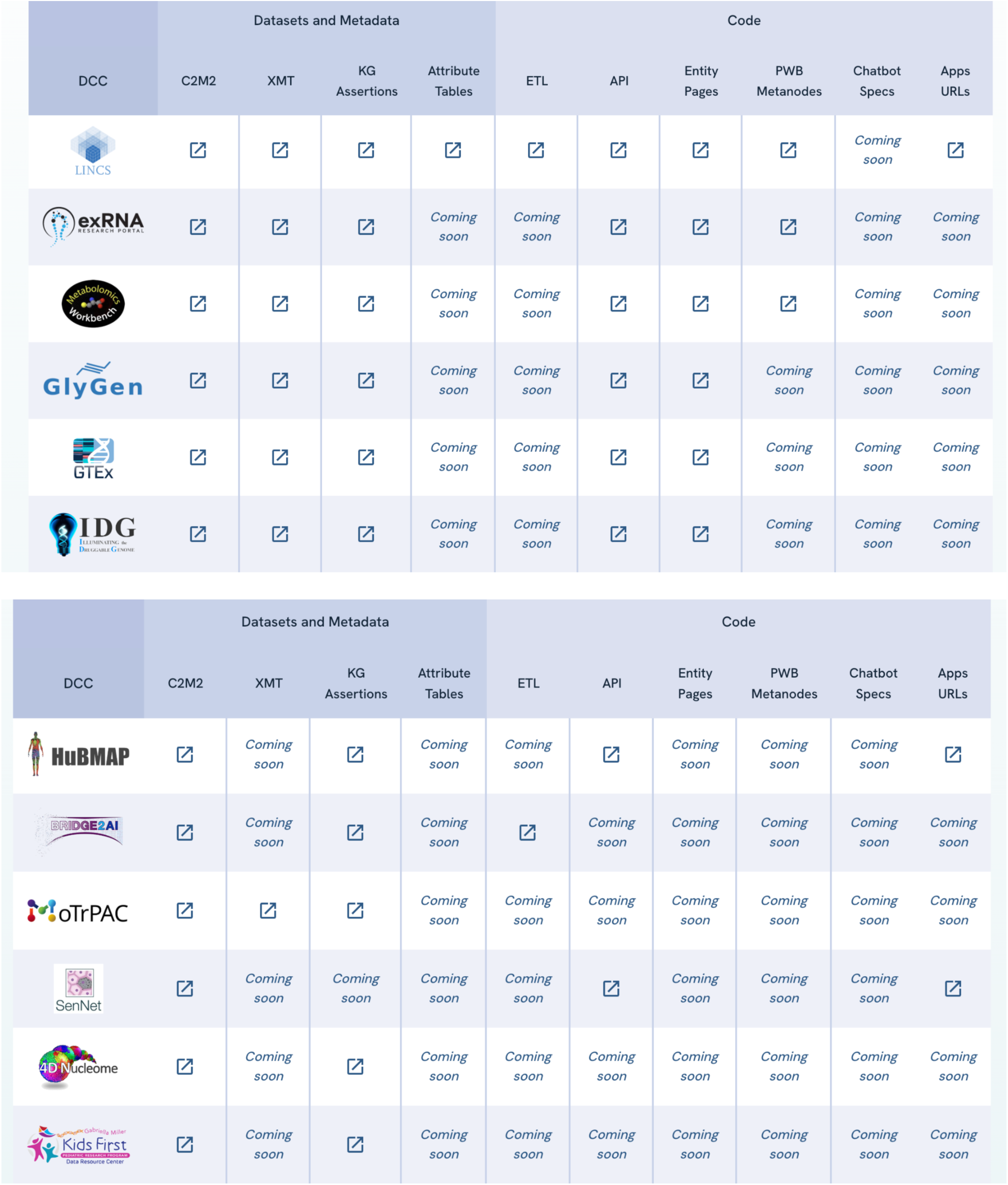
Screenshot from the CFDE Workbench Data Matrix.

### The CFDE Workbench Data Portal Search Engine

The CFDE Workbench Data Portal search engine was developed to enable user-friendly queries of CF programs’ highly processed data, metadata, CF-generated tools, and analysis platforms developed by the CFDE and by the CF programs’ DCCs (https://data.cfde.cloud/). To expand the FAIRness of CF-generated resources, the CFDE Workbench search engine serves the ingested and digested C2M2 packages [20] and several other processed data formats including XMTs, Knowledge Graph (KG) assertions, and attribute tables, all from one search interface. These processed data formats make information derived from the underlying CF programs readily accessible with the aim of easier cross-DCC data integration. Besides serving data and metadata, the search results are linked to downstream data analysis bioinformatics tools. The CFDE Workbench search engine supports search syntax. Entering two search terms together (term1 term2) works as an AND operator, while adding the word OR between search terms (term1 or term2) returns the union of searching each term individually. The dash symbol (-term1) can be used to exclude a specific search term and the double quotes (“term1 term2”) preserve the ordering and consider the search as a bigram. The processed data and metadata formats derived from the DCCs are assembled and incorporated into a PostgreSQL database through a series of scripts. Assets are registered to dedicated tables and linked to a common table indexed for full-text search when applicable. The C2M2 metadata standard describes experimental data across the CF DCCs, linking biosamples and files together with standardized ontologies [31]. The C2M2 metadata is ingested into the PostgreSQL database, and several metadata views are made searchable, for example, raw data files cataloged by the DCCs. The PostgreSQL function websearch_to_tsquery is used in the query to support the search syntax. An XMT file is a sparse matrix serialization format for set libraries where the X refers to the elements in the sets. For example, a Gene Matrix Transpose (GMT) is a library of gene sets, and a Drug Matrix Transpose (DMT) is a library of drug sets. Searchable terms that describe sets are stored in the database and linked to the actual set of entities defined in the XMT record. KG assertions are triples containing a single assertion that defines a typed relationship between two typed nodes from the Data Distillery (DD) KG. Searchable typed nodes from the KG assertions and typed relationships are stored in the database, and these assertions are linked to a record dedicated to the assertion. Where applicable, records are associated with the source data file and the source DCC. The PostgreSQL database is queried by a Next.js webserver that provides an interface for listing DCC assets. Searching for assets is based on their description and other content found within the processed data assets, and it is filtered for asset type or DCC. A search query returns C2M2 metadata (Fig. 4A) and processed data search results (Fig. 4B) respectively. C2M2 metadata search returns projects based on the query term. Users can further refine the results by disease, species, anatomy, gene, protein, compound, data type, assay type, and DCC. The whole or selected query result can be downloaded in JSON format containing the project details and metadata information. The respective project page, also referred to as the records page since additional context such as species, anatomy and disease are specified, provides related biosamples, subjects, collections, and files, some with persistent IDs and access URLs. Some or part of this information can be downloaded in JSON format. The processed data search returns results based on the entity type. The entity types include genes, gene sets, diseases, anatomies, proteins, phenotypes, and more. From the entity pages, the CFDE Workbench provides CFDE-developed tools to help users further investigate knowledge about the entity of interest. For example, when a user is interested in the gene PFKL (Fig. 4C), four CFDE tools that can be used for the gene analysis are displayed. Pressing the submit button will redirect the user to the tool, and the tool will be invoked with the gene or gene set, data table, descriptions, and other items returned from the search.

**Fig. 4.**
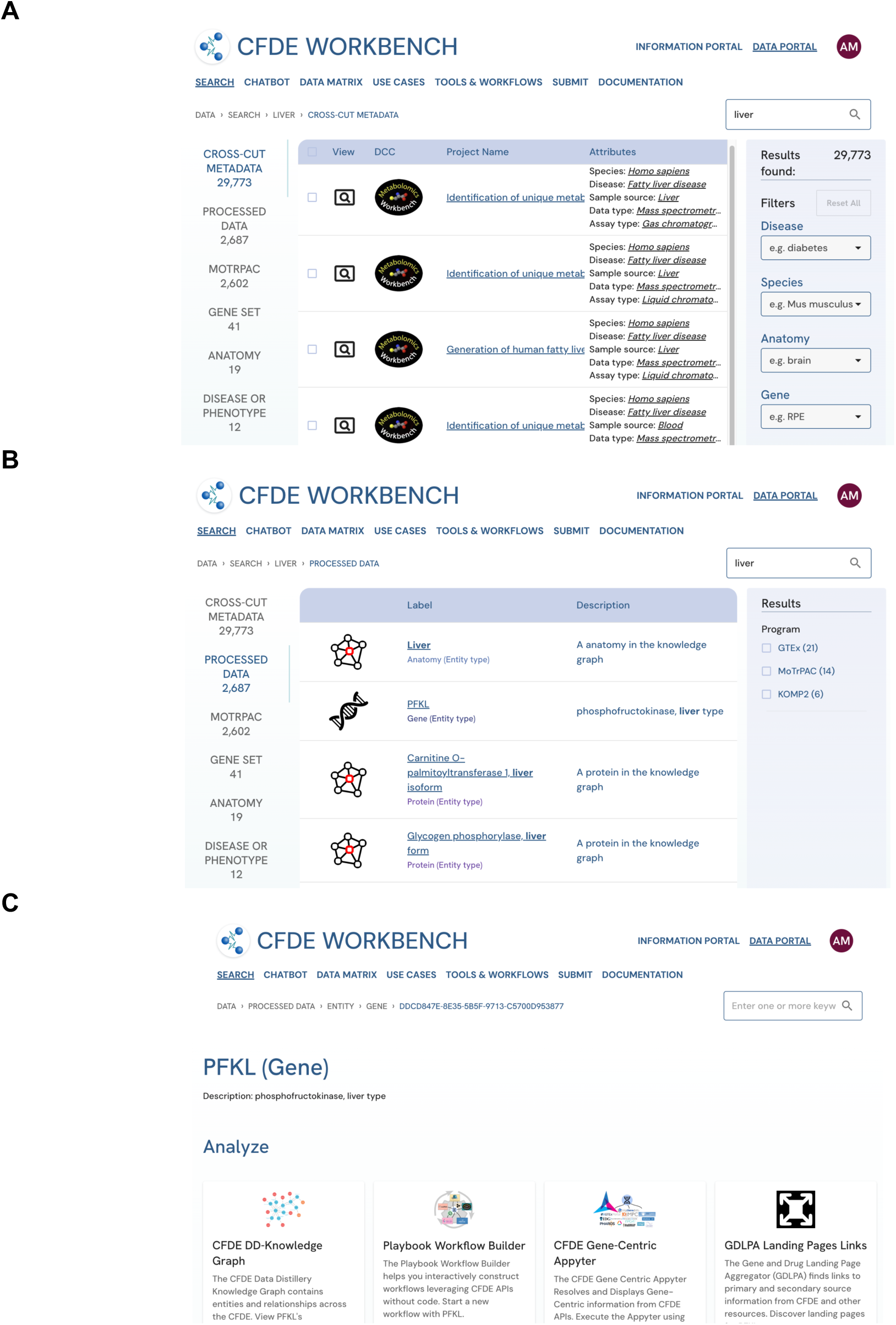
Screenshot from the CFDE Workbench search engine. A) Results from the C2M2 search for the search term “liver”. B) Results from the processed datasets search for the search term “liver”. C) Gene landing page for the gene PFKL.

### The CFDE Workbench Chatbot

The advent of large language models (LLMs) presents the opportunity to enhance the user experience of CFDE Workbench users by implementing a chatbot. We have implemented such a chatbot for the CFDE Workbench (https://data.cfde.cloud/chat) (Fig. 5). The chatbot can answer questions about CF programs and CFDE Data Coordinating Centers (DCCs) and execute queries to retrieve information and data from DCC APIs. By building an interface between the Playbook Workflow Builder (PWB) [32] and the chatbot can access standardized workflows that draw knowledge from multiple DCCs. The PWB provides a simple mechanism for exporting workflows into a programmatic query which can then be executed through the PWB API. These workflows are added to the chatbot to enable users to run processes directly in the chat. A collection of APIs, from the PWB or other relevant APIs from CFDE DCCs, along with robust descriptions of their function and returned results constitutes “Chatbot Specs”. Additionally, the OpenAI ‘assistants’ API enables accurate, responsible, and reproducible results through two mechanisms: 1) Function calls that utilize the Chatbot Specs submitted by DCCs; and 2) Attached files that can be referenced by the chatbot and are useful for citing specific aspects of DCCs that are described in a text document (Fig. 6). Overall, the CFDE chatbot helper provides users with an alternative useful way to obtain information about CF programs, and to perform analyses using CF datasets and API. The chatbot uses APIs to retrieve DCC specific information, mitigating ‘hallucinations’ and providing useful functionality to CFDE Workbench users. The OpenAI Assistants module (https://platform.openai.com/docs/assistants) performs function calling and information retrieval using a ChatGPT model. Information retrieval can be performed on any attached document. In this case, a text file containing information about the CFDE and the DCCs is provided to the API workspace. A system prompt provides directions to strictly use information from this file in answering queries concerning any CF program. Additionally, the assistant is instructed only to answer questions related to the CFDE, the CF, and the DCCs. Any relevant information the assistant should be able to answer can be added to the attached document to provide accurate responses to users. To perform data queries, the assistant can access a collection of function definitions representing discrete PWB workflows. When one of these functions is selected by the assistant, it collects any relevant and required parameters from the user’s query, and then the workflow executes via the PWB API. Frontend components adapted from the PWB metanodes are then used to visualize the results returned from the functions invoked via the chat. In addition to providing such visualizations directly in the chatbot interface, the interface provides a link to executed PWB workflows in the PWB interface for modifications and expansion. While the assistant remains the same for all users, when a user begins a chat, a new thread is created so that the model knows about the conversation history with the individual user. This enables that follow-up questions consider prior context for improved user experience.

**Fig. 5.**
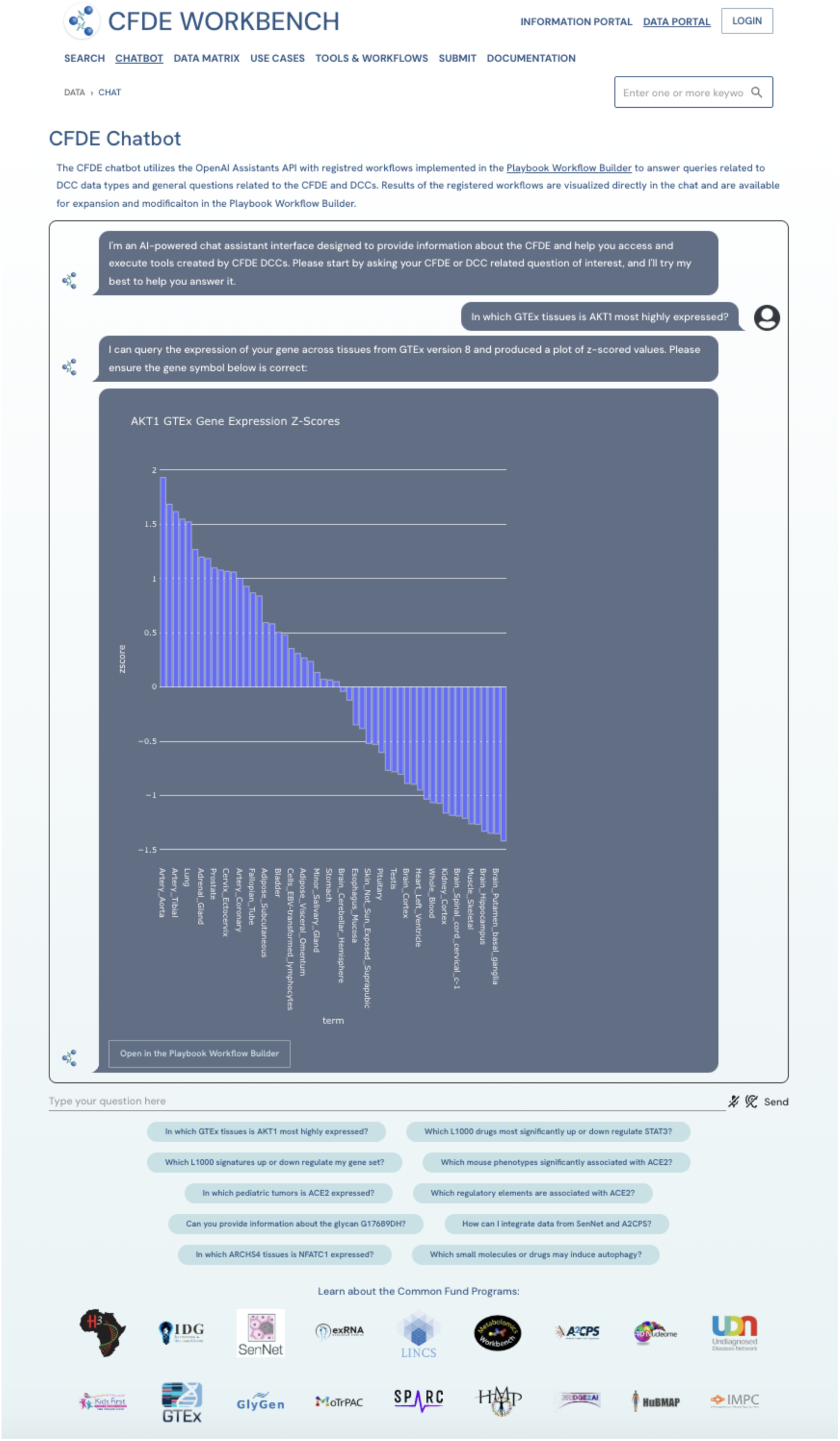
Screenshot from the CFDE Workbench chatbot interface. The results shown are from asking the chatbot to provide the expression levels of the gene ACE2 across human tissues using the GTEx API.

**Fig. 6.**
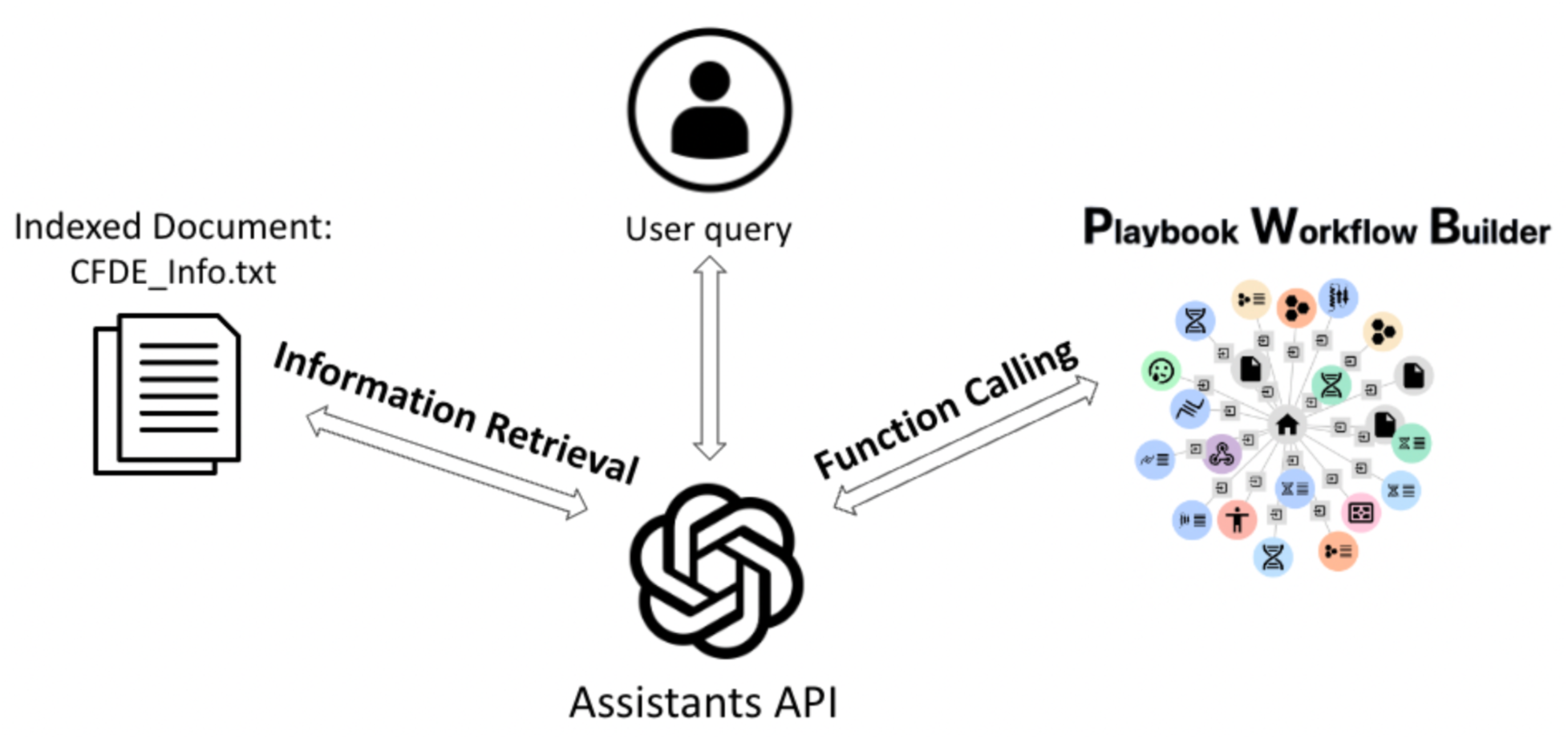
The CFDE Workbench chatbot uses documents describing information about the CF, CF programs, and CF programs’ DCC, as well as the PWB API to answer complex questions and interact with users via a chat interface.

### CFDE Workbench Downstream Analysis Tools

Organizing the data, metadata, and APIs from a collection of CF programs facilitates downstream analysis tools that draw from these resources to enable hypothesis generation that combine CF datasets and tools from multiple programs. So far, these downstream analysis tools and platforms include PWB [32], GeneSetCart (https://genesetcart.cfde.cloud/), CFDE Gene Set Enrichment (CFDE-GSE) (https://gse.cfde.cloud/), Gene and Drug Landing Page Aggregator (GDLPA) [33], FAIRshake [21], and CFDE-KG-UI (https://dd-kg-ui.cfde.cloud/). To demonstrate the utility of the PWB, GeneSetCart, CFDE-GSE, and CFDE-KG-UI the CFDE Workbench has a dedicated section that provides linkable cards to use cases created with these downstream tools and platforms (https://data.cfde.cloud/usecases). The PWB is a web-based platform that facilitates knowledge resolution by enabling users to utilize an ever-growing network of input datasets, semantically annotated API endpoints, and data visualization tools contributed by members of the CFDE. The web-based user-friendly interface enables users to construct workflows using DCC-contributed building-blocks without technical expertise. The output of each step of a workflow is provided in a report containing textual descriptions, interactive and downloadable figures and tables, and an abstract combining the descriptions of each step. GDLPA has links to 53 gene, 18 variant, and 19 drug landing webpages that host gene, variant, and drug landing pages including those produced by CF DCCs. Users can search by gene, variant, or drug terms and the sites that contain knowledge about the gene, variant, and drug are displayed as clickable cards. The FAIRshake platform can be used to assess the level of FAIRness of digital objects by matching each object with a set of questions (FAIR metrics). The collection of answers to these questions is visualized as an interactive insignia. The CFDE knowledge graph user interface (CFDE-KG-UI) is a web interface that interacts with data collected for a CFDE partnership project called the Data Distillery. The CFDE-KG-UI leverages the Cytoscape network visualization library to visualize queries against the knowledge graph database that is stored in a Neo4j database. The other two CFDE Workbench downstream analysis tools, CFDE-GSE and GeneSetCart, are described in the following two sections.

### The CFDE Workbench Gene Set Enrichment (CFDE-GSE) Tool

To further enable discovery across CF programs, the DRC systematically processed CF datasets into gene sets. These gene sets are served from the Data Matrix via another downstream tool called CFDE-GSE. CFDE-GSE stores gene set libraries created from CF programs’ datasets in a Neo4j knowledge graph database. CFDE-GSE then hosts these libraries via a web-based application that serves these gene sets for integrated enrichment analysis. So far, 10 gene set libraries have been created by processing datasets from eight CF programs: LINCS [11], GTEx [1], Metabolomics [5], IDG [10], GlyGen [6], KOMP2 [4], MoTrPAC [18], and HuBMAP [33]. To use CFDE-GSE, gene sets obtained from omics experiments by individual investigators can be submitted to the tool using a simple input form (Fig. 7). The resulting subgraph produced by CFDE-GSE illuminates connections between the input gene set and various CF gene sets that overlap with the queried gene set. Such analysis enables users to explore CF datasets to further hypothesis generation. Besides enrichment analysis, CFDE-GSE can be used to query the consolidated CF knowledge by searching for cross-CF-dataset-associations for a gene or a biological term. Users can query the tool to find cross-DCC information about a drug, a metabolite, a tissue, a cell type, or a phenotype. This feature of CFDE-GSE can be used to discover connections between entities that may not be obvious. Future releases of CFDE-GSE would extend the collection of gene sets to serve data from more CF programs as well as add predicted associations with graph-based machine learning methods, opening the door to a broader cross CF knowledge discovery.

**Fig. 7.**
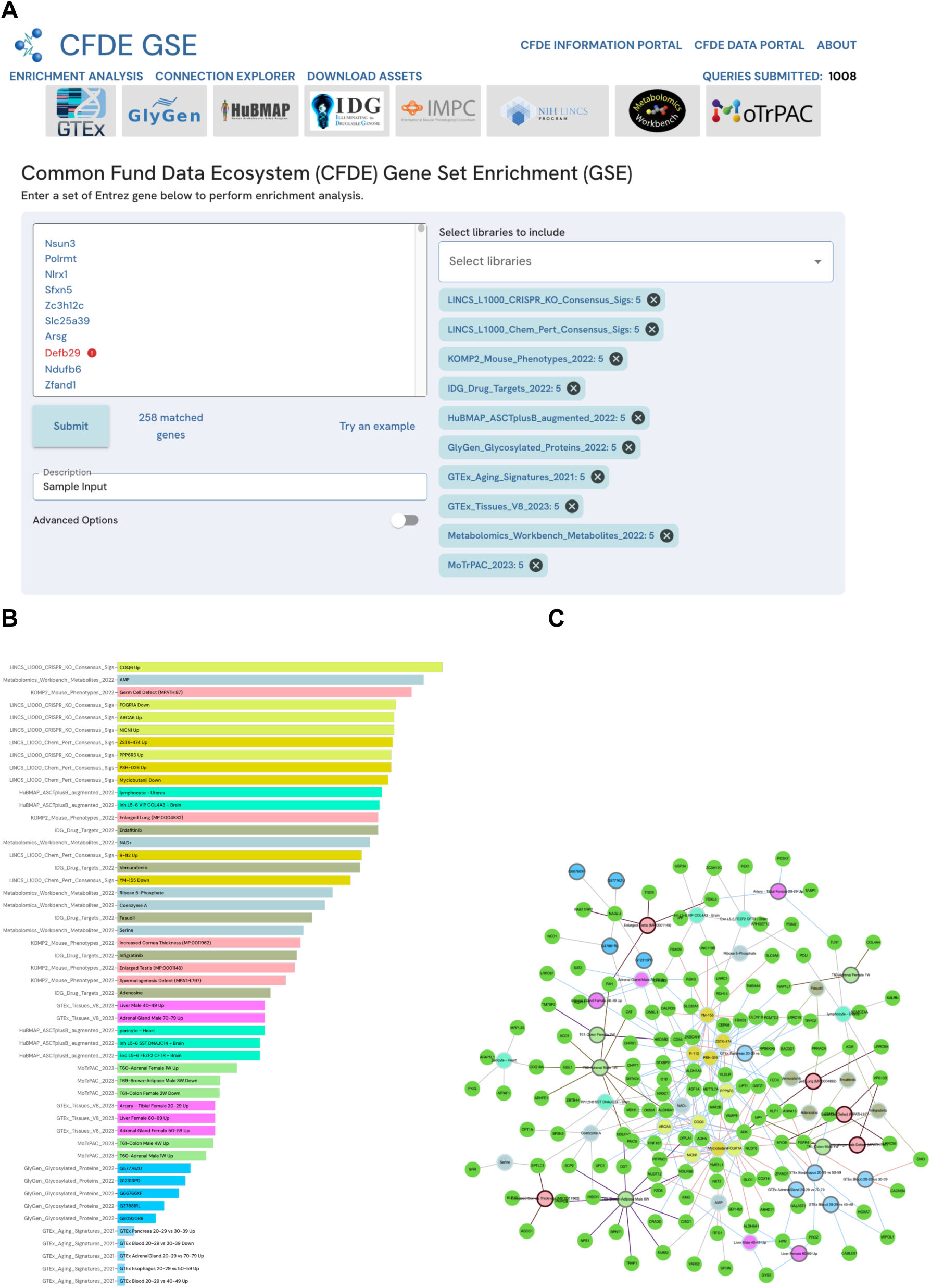
Screenshots from the CFDE tool CFDE-GSE. A) Using the default example gene set and selecting all available gene sets from all DCCs. B) Enrichment results displayed as a bar chart. C) Enrichment results displayed as a ball-and-stick diagram.

### The CFDE Workbench GeneSetCart Platform

To facilitate data integration, discovery, and hypothesis generation across diverse CF programs, highly processed data can be converted into gene, metabolite, drug, and other set libraries. Such abstracted versions of the data facilitate data integration and discovery. GeneSetCart is an interactive web-based platform that enables users to fetch gene sets from various CF programs, augment these sets with gene-gene co-expression correlations or protein-protein interactions, perform set operations such as union, consensus, and intersection on multiple sets, and visualize and analyze these gene sets from one place in a cloud user workspace. GeneSetCart provides access to the same gene sets provided by CFDE-GSE via a term query interface that returns gene sets related to biomedical terms sourced from the CF processed datasets that are hosted by the CFDE Workbench. GeneSetCart also supports the upload of single and multiple user-generated gene sets. Users of GeneSetCart can also obtain gene sets by searching PubMed for genes co-mentioned with any search term. Visualizations of gene sets from various sources can be generated in the form of publication-ready Venn diagrams, heatmaps, UMAP plots, SuperVenn diagrams, and UpSet plots. These visualizations summarize the overlap and similarity between selected sets. Users can also perform functional enrichment analysis on their assembled gene sets through external links to gene set analysis tools such as CFDE-GSE, SigCom LINCS [34], Enrichr [35], Rummagene [36], RummaGEO [37], Kinase Enrichment Analysis (KEA3) [38], and Chip-X Enrichment Analysis (ChEA3) [39]. All gene sets added or created in a session can be saved to a user account for re-analysis and reproducibility. Importantly, GeneSetCart has a feature that enables users to cross gene sets created from data collected by different CF programs (Fig. 8). Overall, GeneSetCart is a useful resource to facilitate hypothesis generation across CF datasets and programs.

**Fig. 8.**
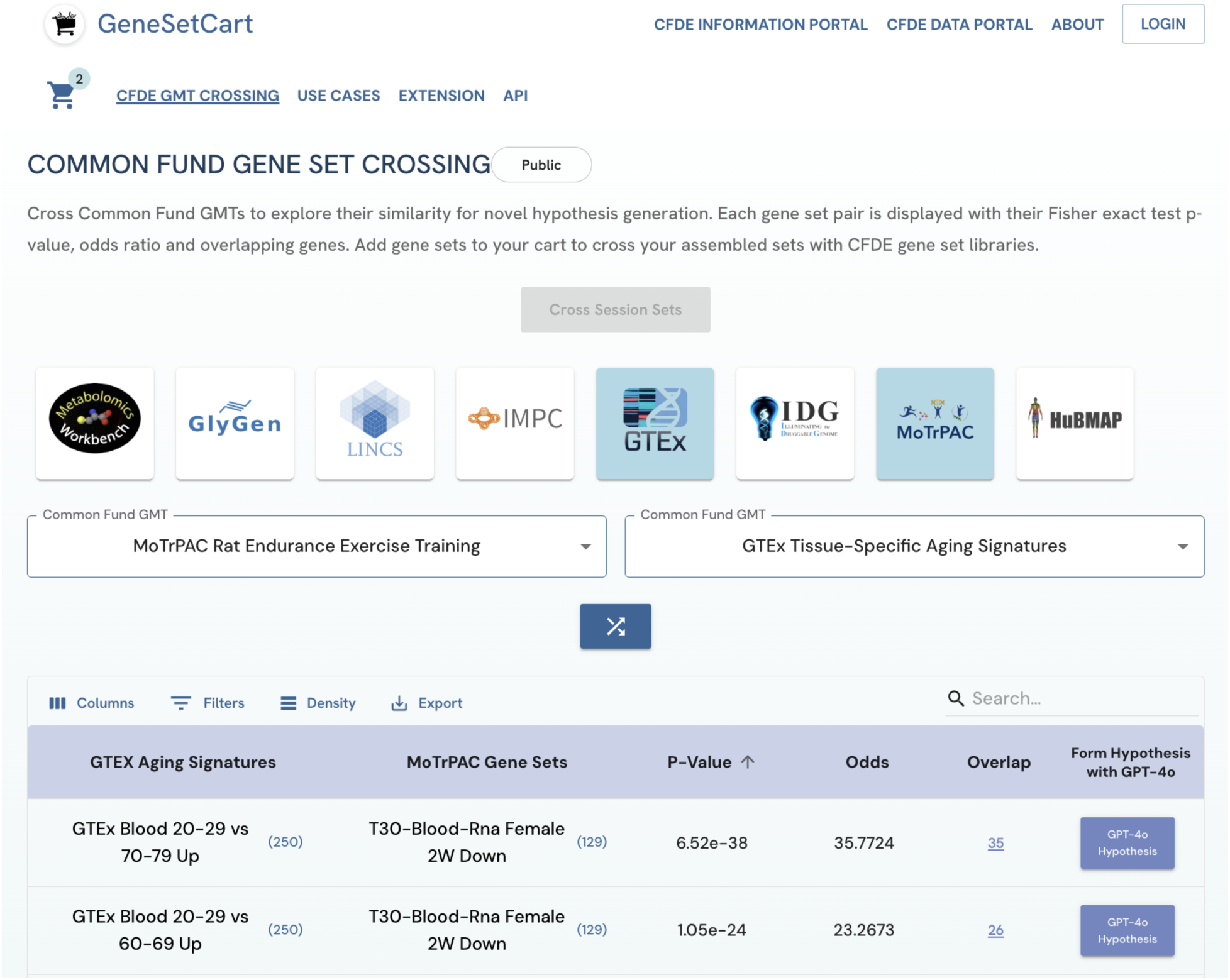
Screenshot from the GeneSetCart platform’s feature that enables users to cross gene sets created from different CF programs. In this example, aging gene sets created from GTEx are crossed with exercise gene sets created from MoTrPAC. The gene sets from GTEx are differentially expressed genes when comparing young (20-29 yrs) to old samples (30-39, 40-49, 50-59, 60-69, and 70-79 yrs). The gene sets from MoTrPAC are differentially expressed genes before and after rats were subjected to an aerobic exercise program. Both datasets were generated with RNA-seq applied to postmortem tissues.

### The CFDE Workbench Cloud Infrastructure

The CFDE Workbench infrastructure is hosted within an Amazon Web Services (AWS) cloud account with different components of the workbench, for example, the Information Portal, the Data Portal and the chatbot, served by different Docker containers orchestrated by Kubernetes [40].

## CONCLUSIONS AND FUTURE DIRECTIONS

By combining the data and information portals (Tables 2-3), the CFDE Workbench is a comprehensive resource where users can collect both information and data from CFDE and CF resources, as well as query disease, gene, drug, and other biological entities across standardized data formats from each CF DCC. For example, for C2M2, the search along with filters may support cohort selection for biomedical researchers. Most importantly, as the CFDE Workbench is expected to continue to evolve over the next few years, the CFDE Workbench interface will be extended to support new datasets and tools, as well as integrate information and knowledge with the other four CFDE centers, making it a valuable resource for promoting FAIR data analyses across CF programs. There are several specific areas where the CFDE Workbench can be improved. Within the Data Portal, AI readiness can be improved by converting data into data frames that can be easily loaded into AI workflows and by making the data accessible via API to enable chatbots and other applications to utilize these APIs. The C2M2 has limited ability to capture patient metadata from CF programs such as GTEx, Kids First, and Bridge2AI. For researchers who work with protected health information (PHI), developing operability with databases such as the database for Genotypes and Phenotypes (dbGaP) [41] and platform such as CAVATICA [42] can enhance the utility of the workbench. We intend to expand the C2M2 to include patient metadata such as body mass index (BMI), smoking status, and other non-identifying health information. The Data Portal’s “Tools and Workflows” page could be improved by adding more linked analysis tools and providing a directory to guide users in selecting the most applicable tools for their data. Lastly, converting C2M2 data from its current relational database to a KG database, such as Neo4j, may provide a better fit for the current metadata model and improve search performance; this effort is already underway. Within the Information Portal, possible areas of improvement include increasing collaboration with similar programs, producing micro-publications that detail each use case, and expanding the submission system to enable DCCs and the CFDE centers to submit events, publications, and use cases directly to the portal.

**Table 2.** Pages that are hosted by the CFDE Workbench Information Portal.

| Portal | Feature | Description | URL |
| --- | --- | --- | --- |
| Information Portal | Landing Page | The information portal contains links to individual DCCs and CF partnership programs, and information about webinars, training and outreach programs, and CFDE documentation. | https://info.cfde.cloud/ |
|  | CF Programs | The CF Program page provides information about the Data Coordination Centers (DCCs) that are participating in the CFDE. | https://info.cfde.cloud/dcc |
|  | Partnerships | The Partnerships page provides information about past and present CFDE partnerships. This includes descriptions, participating DCCs, and related links to publications and other products of the partnerships. | https://info.cfde.cloud/partnerships |
|  | Training & Outreach | The training and outreach events hosted by CFDE centers and DCCs are provided. Filters are available to help find relevant events, and CF community members can request to add events using a form. | https://info.cfde.cloud/training_and_outreach |
|  | Publications | The publications under CF grants are provided along with associated DCC tags, links, and downloadable citations. | https://info.cfde.cloud/publications |
|  | Webinars | The Webinars page features CFDE upcoming and past webinars with their title, presenters, summaries, and video links. | https://info.cfde.cloud/training_and_outreach/cfde-webinar-series |
|  | Documentation | The Documentation page contains descriptions of CFDE consortium standards, protocols, user guides, and other documents generated by the CFDE centers, partnerships, and DCCs. | https://info.cfde.cloud/documentation |
|  | What's New? | The latest updates to the portal include improvements and newly added assets and events. | https://info.cfde.cloud/news |

**Table 3.**
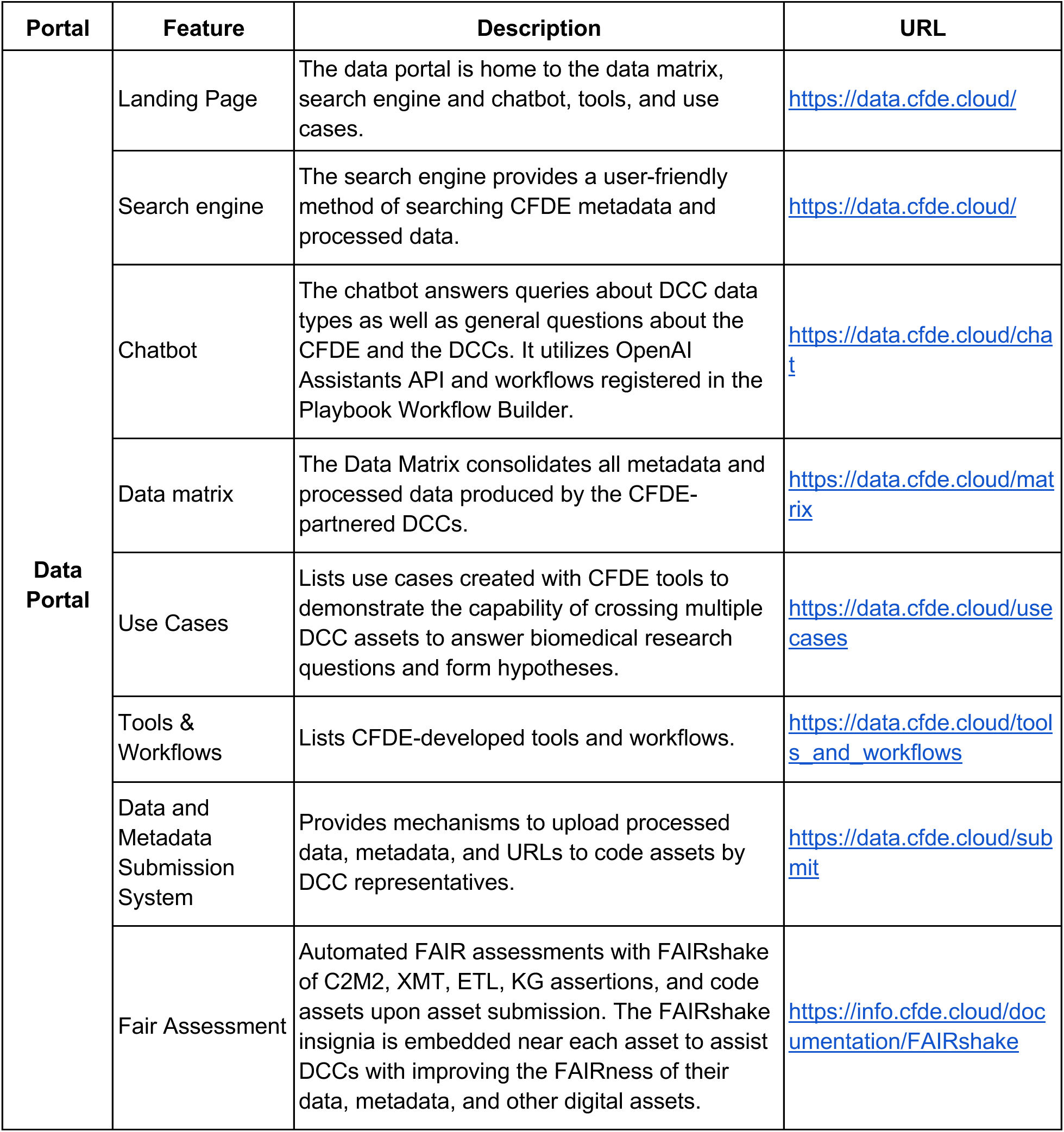
Pages that are hosted by CFDE Workbench Data Portal.

| Portal | Feature | Description | URL |
| --- | --- | --- | --- |
| Data Portal | Landing Page | The data portal is home to the data matrix, search engine and chatbot, tools, and use cases. | https://data.cfde.cloud/ |
|  | Search engine | The search engine provides a user-friendly method of searching CFDE metadata and processed data. | https://data.cfde.cloud/ |
|  | Chatbot | The chatbot answers queries about DCC data types as well as general questions about the CFDE and the DCCs. It utilizes OpenAI Assistants API and workflows registered in the Playbook Workflow Builder. | https://data.cfde.cloud/chat |
|  | Data matrix | The Data Matrix consolidates all metadata and processed data produced by the CFDE-partnered DCCs. | https://data.cfde.cloud/matrix |
|  | Use Cases | Lists use cases created with CFDE tools to demonstrate the capability of crossing multiple DCC assets to answer biomedical research questions and form hypotheses. | https://data.cfde.cloud/usecases |
|  | Tools & Workflows | Lists CFDE-developed tools and workflows. | https://data.cfde.cloud/tools_and_workflows |
|  | Data and Metadata Submission System | Provides mechanisms to upload processed data, metadata, and URLs to code assets by DCC representatives. | https://data.cfde.cloud/submit |
|  | Fair Assessment | Automated FAIR assessments with FAIRshake of C2M2, XMT, ETL, KG assertions, and code assets upon asset submission. The FAIRshake insignia is embedded near each asset to assist DCCs with improving the FAIRness of their data, metadata, and other digital assets. | https://info.cfde.cloud/documentation/FAIRshake |

## ACKNOWLEDGEMENTS

This project was funded by NIH grants OT2OD036435 (CFDE Workbench). We would like to give special thanks for Drs. Kristin Kano, Karlie R. Sharma, Haluk Resat, George J. Papanicolaou, and Chris Kinsinger for useful feedback. We are also grateful for the web design team from Indicius Inc. for their contribution.

